# Neurophysiological network dynamics of pitch change detection

**DOI:** 10.1101/2020.10.07.328450

**Authors:** Soheila Samiee, Dominique Vuvan, Esther Florin, Philippe Albouy, Isabelle Peretz, Sylvain Baillet

## Abstract

The detection of pitch changes is crucial to sound localization, music appreciation and speech comprehension, yet the brain network oscillatory dynamics involved remain unclear. We used time-resolved cortical imaging in a pitch change detection task. Tone sequences were presented to both typical listeners and participants affected with congenital amusia, as a model of altered pitch change perception.

Our data show that tone sequences entrained slow (2-4 Hz) oscillations in the auditory cortex and inferior frontal gyrus, at the pace of tone presentations. Inter-regional signaling at this slow pace was directed from auditory cortex towards the inferior frontal gyrus and motor cortex. Bursts of faster (15-35Hz) oscillations were also generated in these regions, with directed influence from the motor cortex. These faster components occurred precisely at the expected latencies of each tone in a sequence, yielding a form of local phase-amplitude coupling with slower concurrent activity. The intensity of this coupling peaked dynamically at the moment of anticipated pitch changes.

We clarify the mechanistic relevance of these observations in relation to behavior as, by task design, typical listeners outperformed amusic participants. Compared to typical listeners, inter-regional slow signaling toward motor and inferior frontal cortices was depressed in amusia. Also, the auditory cortex of amusic participants over-expressed tonic, fast-slow phase-amplitude coupling, pointing at a possible misalignment between stimulus encoding and internal predictive signaling. Our study provides novel insight into the functional architecture of polyrhythmic brain activity in auditory perception and emphasizes active, network processes involving the motor system in sensory integration.

## Introduction

Pitch is a fundamental perceptual feature of sound that is a form-bearing dimension of music and an important cue for understanding speech (Bregmann, 1990). Meaningful pitch changes are perceived by combining auditory inputs with contextual priors and by engaging attentional focus (Garrido et al., 2007). Hence, pitch perception engages a network of brain regions also involved in auditory prediction and perceptual decision making (Peretz and Zatorre 2005). Here, we sought to determine the regional and network neurophysiological mechanisms and dynamics crucially involved in brain pitch processing.

One classic approach to assess the neural processing of pitch is via the oddball paradigm (Näätänen et al., 2007), which consists of the presentation of sequences of identical tones that are infrequently interrupted by a deviant stimulus that differs in pitch. Electroencephalographic (EEG) and magnetoencephalographic (MEG) oddball studies have shown that when attention is directed away from auditory stimuli, the brain response to deviant pitch is marked by an early negative event-related component (Mismatch Negativity, Näätänen et al., 2007). Later positive components, the P3a and P3b, reflect the orientation of attention towards the deviant sound and are typically elicited by the detection and evaluation of the deviant target. Anatomically, the P3a is generated by frontal attention-orienting and novelty-processing mechanisms; whereas the P3b is observed when the stimulus is task-relevant and is therefore thought to originate from temporo-parietal brain activity associated with attention (Polich, 2007, see also Opitz et al., 2002; Schönwiesner et al., 2007 for fMRI studies).

However, little is known about the key neurophysiological mechanisms of regional and network brain activity for pitch detection. One way of investigating this question is to contrast the brain activity of typical listeners and participants affected by a pitch-specific neurodevelopmental disorder like congenital amusia (Peretz 2016). Congenital amusia is characterized by a disturbance in the processing of pitch, as documented in oddball (Hyde & Peretz, 2004; Peretz et al., 2005) and delayed match to sample tasks (see Tillmann et al., 2016 for review). Amusic individuals are impaired at detecting pitch deviations that are smaller than two semitones. Specifically, the amusic brain produces a normal mismatch negativity (MMN) response to deviations as small as an eighth of a tone (25 cents) from repeated standard tones, but without conscious perception (Omigie et al., 2012). Furthermore, this MMN is not followed by a normal and typical positivity (P3b) associated with conscious sensory detection (Moreau et al., 2013). This functional gap between non-conscious and conscious pitch change detection in the amusic brain is currently attributed to altered brain connectivity between the superior temporal gyrus (STG) and the inferior frontal gyrus (IFG; Peretz, 2016). Dynamical causal modelling in MEG showed that amusic participants express both decreased backward connectivity between the right IFG and the right auditory cortex during pitch *encoding* (Albouy et al., 2013; Albouy et al., 2019) and decreased forward connectivity during pitch *change detection* (interpreted as reduced prediction error signalling; Albouy et al., 2015). In typical listeners, the role of fronto-temporal network connectivity in pitch change detection has been further documented using measures of oscillatory brain dynamics related to sensory input prediction and the relationship between rhythmic responses and entrained oscillations (Phillipes et al. 2015, Tse et al. 2018, Haegens & Zion Golumbic, 2018; Nobre & van Ede, 2018). Regarding amusia, recent studies have only reported power fluctuations in frequency bands of brain activity taken separately, such as gamma (Albouy et al., 2013) and alpha (Tillmann et al., 2016).

We propose a more integrative view of the neurophysiological dynamics that are crucial to pitch-change perception by assessing the time-resolved variations of interdependent brain oscillations and the directional connectivity between key nodes of the fronto-temporal network recruited in pitch processing. Our recent work highlighted the singular role of beta oscillations directed from the motor cortex to auditory regions during attentive pitch processing of tone sequences (Morillon & Baillet, 2017). These findings are in line with the conception that the motor cortex issues predictive beta-band signals in anticipation to sensory inputs (Michalareas et al. 2016; Baillet 2017; Chao et al. 2018). This signalling mechanism is not entirely captured by mere augmentations in beta-band signal strength. Indeed, Change et al. (2018) observed that beta power was actually reduced in the auditory cortex prior to a predictable pitch change. Morillon & Baillet (2017) showed that the crucial signaling mechanisms was actually in the coupling of motor beta oscillations with the slower delta-band fluctuations entrained by auditory inputs.

In the current study, we dissected in further detail the network dynamics that are essential to detecting subtle pitch changes in tone sequences, using data from both typical and amusic participants. We considered the latter as a group model of perturbed pitch perception, in order to highlight the neurophysiological components that are essential to conscious detection of pitch changes.

## Results

### Behaviour

Participants listened to a sequence of five pure tones and were asked to categorize the trial as *standard* or *deviant*, based on the fourth target tone in the sequence (Fig. 1 A, see *Material and Methods* for details). The deviant trials had four levels of difficulty depending on difference in pitch frequency between the fourth tone and the other tones in the sequence presented at a standard pitch (1^st^ to 3^rd^ + 5^th^ tone). The pitch differences used amounted to 25, 50, 100, and 200 cents (percent of a semitone). In standard trials, all tones in the sequences were presented using the same identical pitch. We measured the hit-rate to capture behavioral performance, defined as the ratio of correctly detected trials with respect to all trials for all five deviance levels (the four possible deviant pitch levels and the standard, Fig. 1B). A two-factor (group × deviance level) between-subject ANOVA of observed hit rates revealed a significant interaction between groups and deviance levels (F (1, 4) = 19.1, p < 0.001). Post-hoc analysis indicated no difference in hit-rate between deviance levels of 25 and 50 cents (post-hoc Tukey test, t(70) < 1.1, p_Tukey_ > 0.98), and between deviance levels of 100 and 200 cents (t(70) <= 0.28, p_Tukey_ = 1), but other interactions were significant (p_Tukey_ < 0.01). We therefore combined the trials with deviance levels of 25 and 50 into a single condition (*Small Deviance*), and all trials with deviance levels of 100 and 200 cents into another condition (*Large Deviance*). We used *d’* to measure sensitivity when assessing behavioral performance in both deviance conditions (Fig. 1C). We found an interaction between the level of target pitch deviance and groups (F(1,1)=20.3, *p <* 0.001). Further, both typical listeners and amusics showed higher sensitivity to large deviance (controls: *t*(60) = 4.1, *p*_*Tukey*_ < 0.001, amusics: *t*(60) = 10.5, *p*_*Tukey*_ < 0.001). Typical listeners were more sensitive than amusics in the Small Deviance condition (*t*(60) = 7.5, *p*_*Tukey*_ < 0.001) but not in the Large Deviance condition (*t*(60) = 1.2, *p*_*Tukey*_ = 0.66).

**Figure 1:**
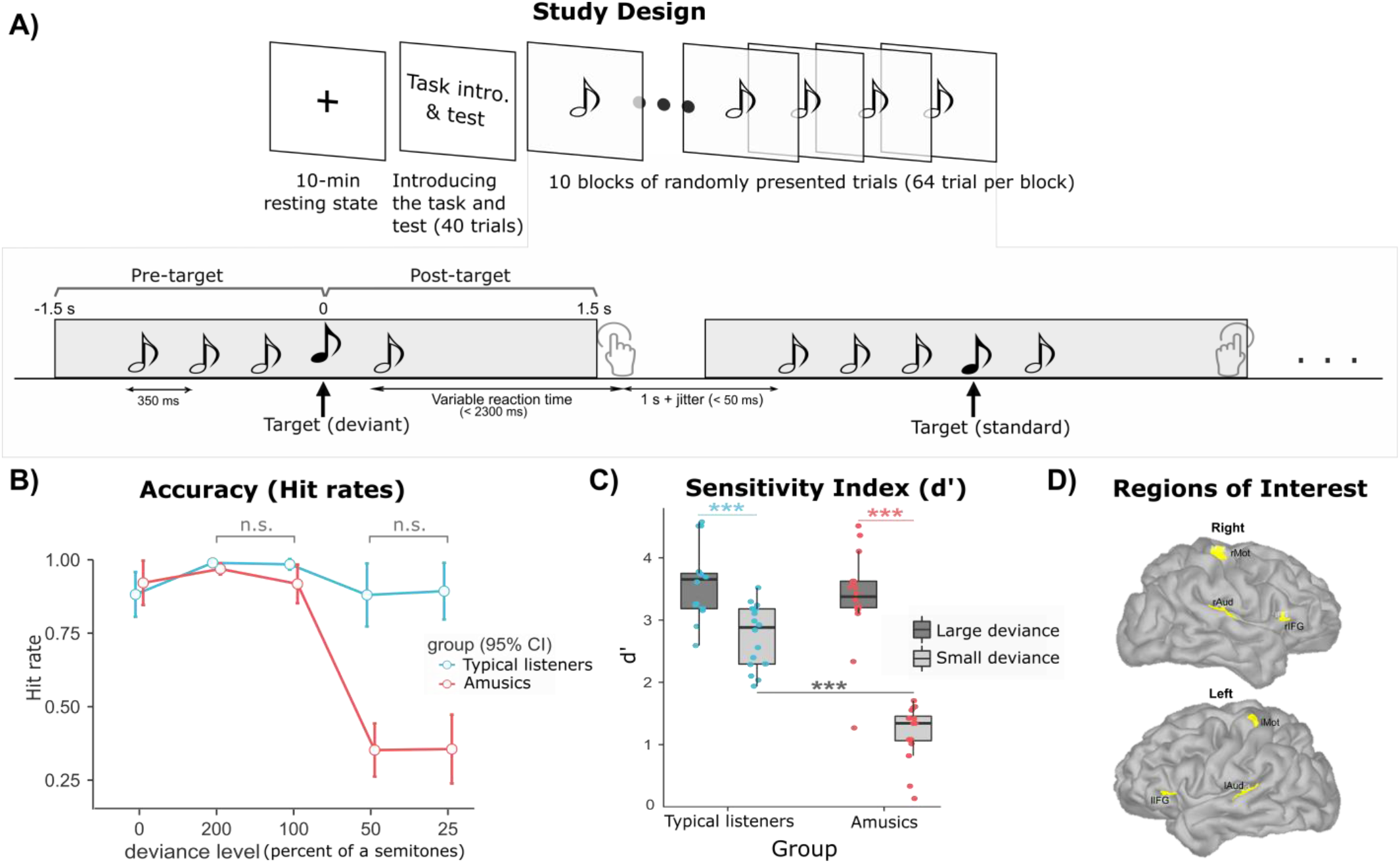
Experimental design, behavioral performance, and regions of interest. A) Study design and nomenclature. B) Hit rate for 5 different deviance levels of target tone (with respect to the first three tones in each trial): 0 deviance reflects the condition with all tones having the same pitch frequency (*standard* trials). In all other four conditions, the fourth tone in the sequence was a deviant pitch tone. There were no significant interactions between group for pitch deviance levels of 25 and 50 cents, and for 100 and 200 cents. C) Sensitivity index (*d’*) for amusic participants and controls in the large and small deviance conditions. D) Anatomical regions of interest shown in yellow on the cortex of a representative subject: These ROIs were identified using functional localizers registered to the anatomy of each participant.

### Coupling between slow and fast neural dynamics

Across participants and for both groups, we observed the strongest phase-amplitude coupling (PAC) for the entire sequence in the right auditory cortex (rAud) between the phase of delta-band activity at [2,4] Hz and the amplitude of neurophysiological signals in the beta frequency range at [15, 35] Hz (Fig. 2A). Time-resolved measures of PAC variations (tPAC) in rAud over the tone sequence are shown in Fig. 2B. The five data points in Fig. 2B report tPAC values during the presentation of each of the five auditory tones. The last two tPAC data points correspond to subsequent time windows during which there was no tone presentation. We found in both groups and across all tested time windows that the strength of phase-amplitude coupling was above chance levels (z > 3.4, p_corrected_ < 0.01). Overall, coupling was stronger in amusics than in typical listeners (F(1)=11.1, p < 0.001), with no effect of response accuracy (F(1)=0.02, p = 0.88) or pitch deviance (F(1)=0.94, p = 0.33; Fig. 2C). There was also a main effect of time (F(6)=6.5, p < 0.001): in both groups, a post-hoc analysis showed that PAC increased after the onset of the tone sequence (p = 0.0006) and decreased after the occurrence of the target tone (over the three subsequent time windows: p = 0.019, p = 0.013, and p < 0.0001, respectively).

**Figure 2:**
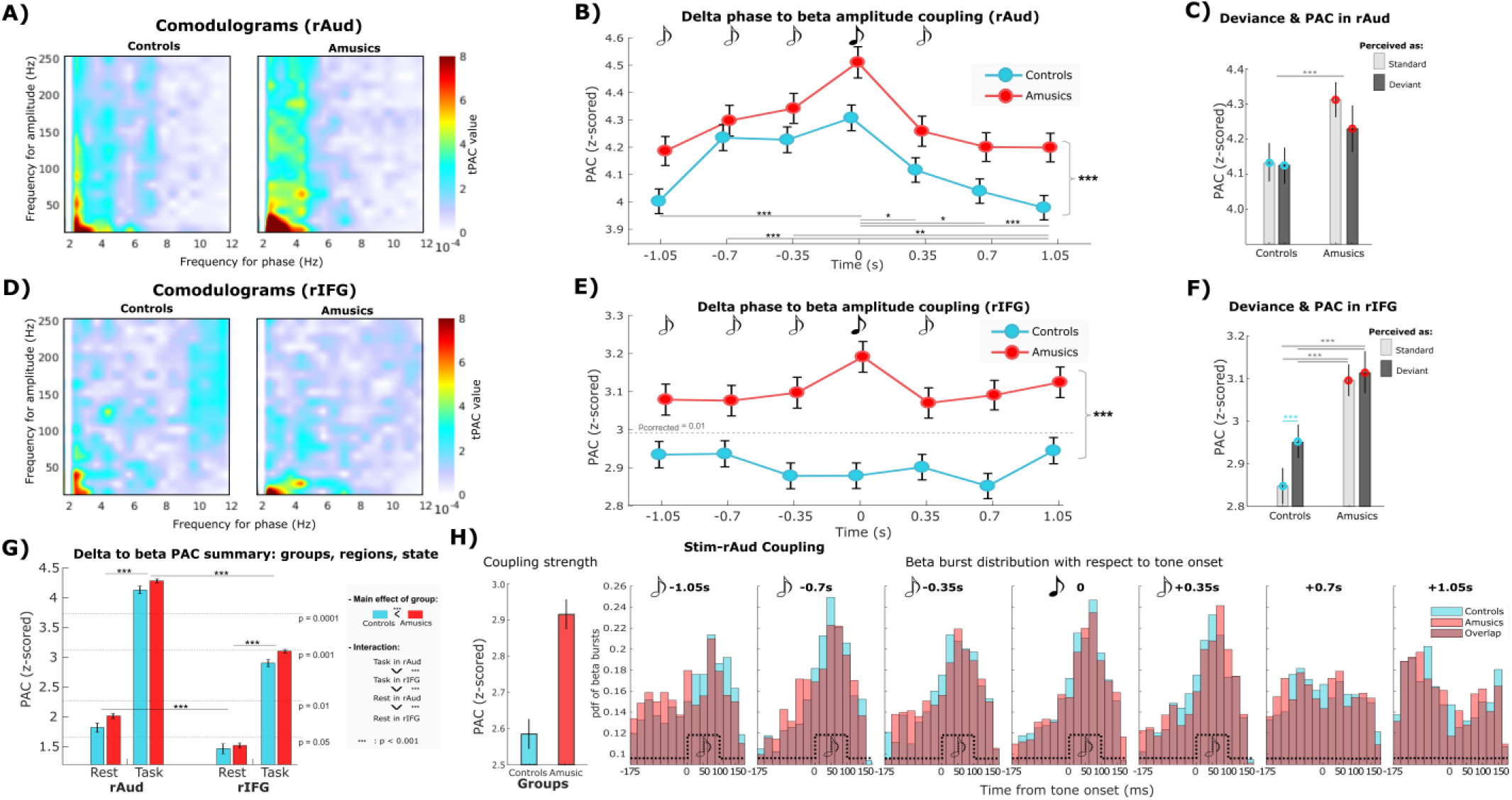
Phase-amplitude coupling analyses. A) The comodulograms extracted from source signals in the rAud ROI are shown for controls and amusics. In both groups, the strongest mode of phase-amplitude coupling was between the phase of delta activity at [2,4] Hz and the amplitude of neurophysiological signals in the beta range at [15,35] Hz. B) Time-resolved analysis of phase-amplitude coupling (tPAC) between these frequency ranges in rAud, z-scored with respect to surrogate data (z = 3.6 was above chance levels with corrected p = 0.001 across time windows). C) Interaction between perceived pitch deviance and group, for delta-to-beta PAC in rAud. D) PAC comodulograms of rIFG source signals in both groups. E) Time-resolved delta-to-beta PAC in rIFG, z-scored with respect to surrogate data. F) Interaction between perceived pitch deviance and group, for delta-to-beta PAC in rIFG. G) Delta-to-beta coupling in rAud and rIFG in both groups, during baseline resting-state and task performance. H) Time-resolved stimulus (pure 2.85-Hz sinusoidal signal) to brain (beta activity in rAud) phase-amplitude coupling; Left panel: coupling strength z-scored with respect to surrogate data (z = 2.44 is above chance levels with corrected p = 0.05 across time windows); right panel: The empirical distributions of the preferred phase angles of tPAC coupling (expressed as latencies with respect to tone onsets) between auditory tone presentations and peaks of beta bursts, across trials and for each time window along the tone sequence. Dashed boxcar shapes indicate the onset and duration of each tone. The bars indicate the 95% confidence interval.

In the right inferior frontal cortex (rIFG), phase-amplitude coupling was also the strongest between the phase of regional delta activity and the amplitude of beta-band fluctuations (Fig. 2D). Time-resolved tPAC analysis in that region revealed a main effect for groups (F(1)=43.95, p < 0.0001; Fig. 2E): as in rAud, amusics expressed stronger PAC levels than controls (p < 0.001, Fig. 2E & F). We also observed a main effect of deviance level (F(1)=5.8, p = 0.0157) and an interaction between actual deviance and accuracy of pitch change detection (F(1,1)=13.1, p < 0.001). Indeed, in controls, phase-amplitude coupling was stronger in rIFG when target tones were perceived as deviant than when reported as standards (p_corrected_ = 0.007; Fig. 2F), regardless of response correctness.

We performed a two-factor ANOVA (group x perceived deviance) of phase-amplitude coupling in rIFG, which confirmed a main effect of group (F(1)=77.82, p < 0.0001) and of perceived deviance (F(1)=7.05, p = 0.0079). In the right auditory cortex, there was only a main effect of group (F(1)=21.35, p < 0.0001) and no effect of perceived deviance (F(1)=2.25, p = 0.13). These observations point at a neurophysiological marker in the inferior frontal cortex of the individual’s perception of the target tone as deviant, regardless of accuracy. There was no such effect in the right auditory region, and it was absent in all tested regions in amusic participants.

We also derived phase-amplitude coupling statistics in the baseline resting state prior to the auditory-testing session, with the objective of evaluating a possible predictive relation with the values observed during task performance (Fig. 2G). Resting-state PAC between ongoing delta and beta was above chance level in rAud for both groups (p < 0.05), but only marginally in rIFG (p > 0.07). We found a main effect of group (amusics stronger, F(1)=13.93, p = 0.0002), region (rAud stronger, F(1)=411.44, p < 0.0001) and behavior (resting-state weaker than task performance, F(1)=2241.1, p < 0.0001), with a significant interaction between region and state (F(1,2)=93.97, p < 0.0001). Post-hoc analysis of the interaction showed that PAC in rAud during task performance was stronger than in rIFG (p < 0.0001) and stronger than during the resting state in rAud (p < 0.0001) in both groups.

### Beta bursts are temporally aligned with tone presentations in a sequence

We also derived measures of phase-amplitude stimulus-to-brain coupling in the right auditory cortex. Our observations reproduced previously reported findings (Fujioka et al., 2012; Cirelli et al., 2014; Chang et al., 2018) of stronger coupling in amusics compared to controls between the phase of a reference sinusoid adjusted to the tone sequence and the amplitude of beta signaling (F(1)=60.5, p < 0.001; Fig. 2H: left panel). There was no significant effect of time (F(6)=1.16, p = 0.32), accuracy (F(1)=2.08, p = 0.14) or pitch deviance (F(1)=0.41, p = 0.53). Overall, neurophysiological delta-to-beta phase-amplitude coupling was stronger than stimulus-to-beta coupling in the tested region (t(119985)=69.45, p < 0.001).

For each trial, we also extracted the latency of beta amplitude bursts with respect to the corresponding tone presentation in the sequence. We found in both groups that after the first tone in the sequence was presented, the amplitude of beta bursts was maximal at the expected latency of auditory inputs reaching the auditory cortex (i.e. about 50ms after tone onset; Fig. 2H: right panel).

### Frequency-specific network interactions

We measured the coherence between all pairs of regions of interest. We observed a main effect of the pair of regions of interest (Aud-IFG presented stronger coherence than Mot-Aud and Mot-IFG, F(2) = 120.29, p < 0.0001), frequency band (delta-band coherence was stronger than beta-band’s, F(1) = 49.7, p < 0.0001), and laterality (with right-hemisphere coherence being stronger than left-hemisphere, F(1) = 17.91, p < 0.0001), but not for group (F(1) = 1.56, p = 0.21) or state (rest, vs pre-target vs post-target, F(2) = 0.03, p = 0.96) (Fig. 3A). We also observed significant interactions between frequency-band and group (F(1,1) = 6.16, p = 0.013) with stronger delta coherence in controls than in amusic participants (p_corrected_ = 0.04).

**Figure 3:**
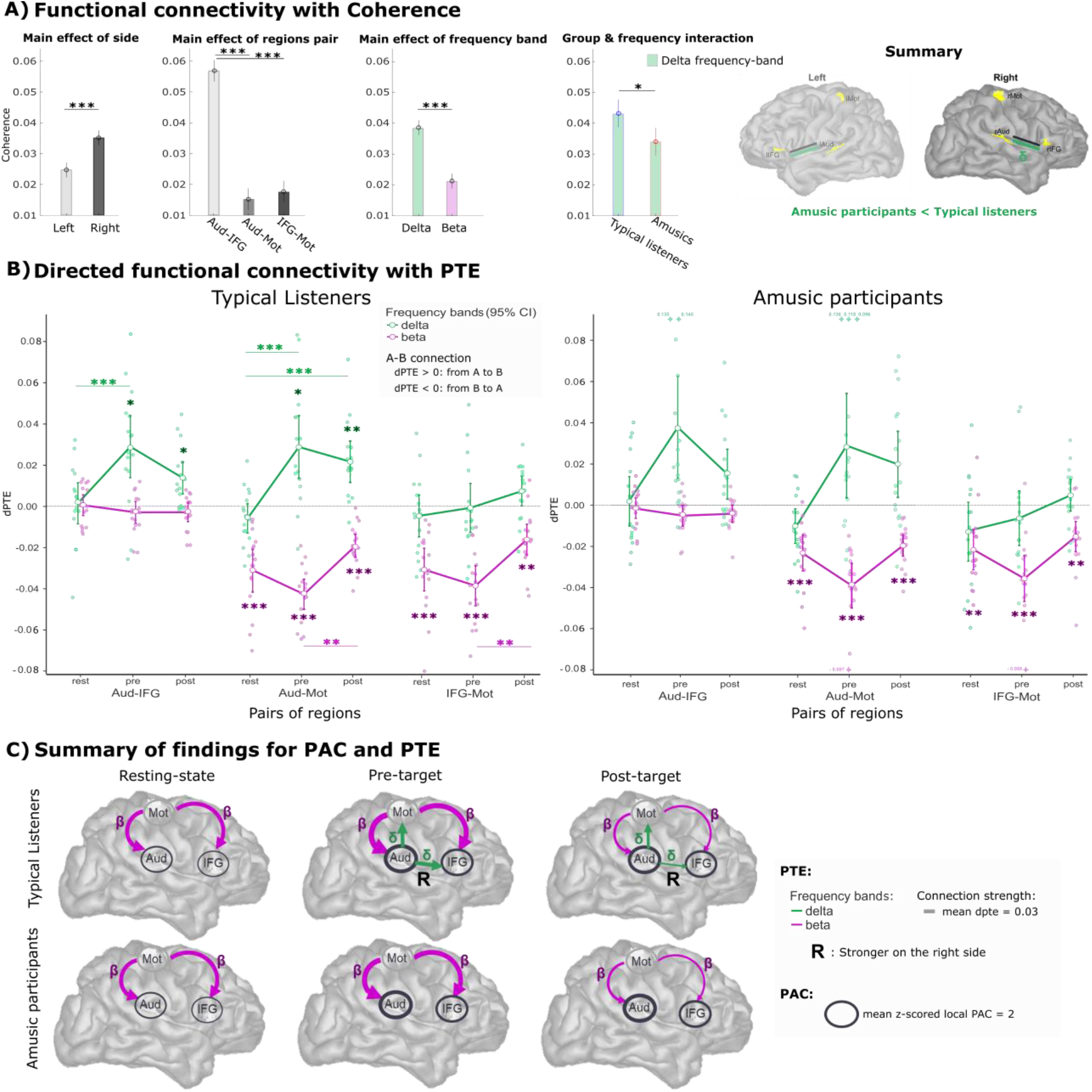
Functional and directed connectivity analyses. A) There was a main effect of laterality in coherence measures, which were stronger between right-hemisphere region pairs; coherence between Aud-IFG was stronger than between the other pairs of ROIs, coherence in the delta band was stronger than in the beta range. There was also an interaction between frequency band and group, with controls expressing stronger delta coherence compared to amusics – *: p < 0.05, ***: p < 0.001 (Tukey’s corrected). These findings are summarized graphically in the right panel. B) Measures of directed phase-transfer entropy (dPTE) for each pair of left and right ROIs, frequency band, and state (resting-state, pre- and post-target tone), for controls (left) and amusics (right). The distributions of dPTE values are shown for each inter-regional connection and over the baseline resting state, the pre- and the post-target segments of the tone sequences. The green and pink traces are for delta- and beta-band dPTE, respectively. The circles show group sample averages, with bars indicating 95 % confidence intervals. All statistics were corrected for multiple comparisons (*: p < 0.05, **: p < 0.01, ***: p < 0.001). C) Graphical summary of findings for PAC and PTE plotted on a template cortical surface: the arrows are schematics for significant dPTE directed connections. Line thickness indicates the strength of dPTE group averages. Similar patterns were found between left-hemisphere ROIs (not shown), with stronger delta-band downstream connectivity from right Aud to right IFG in controls (t(84) = −2.7, p_Tukey_ = 0.04). The thickness of the circles around the auditory (Aud) and inferior frontal (IFG) cortices indicates the strength of local cross-frequency phase-amplitude coupling between the phase of delta-band activity and the amplitude of beta-band activity.

We then measured manifestation of frequency-specific directed interactions between regions of interest, using phase-transfer entropy (PTE; Lobier et al., 2014). The analysis showed that in the resting state of controls, beta-band activity was directed from motor cortex to Aud (t(15) = −6.48, p < 0.001) and from motor cortex to IFG (t(15) = −6.31, p < 0.001; Fig. 3B: left panel). We also found similar expressions of directed connectivity during task performance in both pre-target (Aud: T (15) = −12.42, p < 0.001, IFG: T (15) = −8.42, p < 0.001) and post-target segments (Aud: T (15) = −7.33, p < 0.001, IFG: T (15) = −4.6, p = 0.006). There was a reversed directed connectivity transfer in the delta range, from the auditory cortex to IFG and motor regions in pre-target (IFG: T (15) = 4.06, p = 0.018, Mot: T (15) = 4.07, p = 0.017) and post-target (IFG: T (15) = 3.71, p = 0.038, Mot: T (15) = 4.66, p = 0.006) segments, but not in the resting state.

In typical listeners, a three-factor ANOVA (pairs of ROIs × state × frequency bands) of PTE measures confirmed a significant main effect of the frequency band (F (1) = 4.57, p < 0.001) with opposite directions of connectivity transfer for delta-versus beta-band activity. Interactions showed that directed connectivity transfer of delta-band activity from bilateral auditory regions to inferior frontal cortices was increased during pre-target tone presentations, compared to baseline resting state (t(90) = 4.73, p < 0.001). Delta - band transfer was also stronger from bilateral auditory to motor regions over the entire tone sequence (pre- and post-target) compared to the baseline resting state (pre: F (90) = 5.39, p < 0.001, post: F (90) = 4.27, p < 0.001). Reversed directed connectivity transfers were observed in the beta band from motor to auditory regions, and from motor to inferior frontal cortices. PTE was stronger over the pre-target segment compared to post-target (from motor to auditory regions: F (90) = 3.65, p = 0.006; from motor to inferior frontal cortices: F (90) = 3.52, p = 0.009). All these observations were identical for both hemispheres, with the exception of delta transfer from auditory to inferior frontal cortex, which was stronger on the right side (post-hoc: hemisphere × frequency band interaction: t(84) = −2.7, p_corrected_ = 0.04).

Qualitatively, directed connectivity measures were similar between typical listeners and amusics (Fig. 3B). There was no significant main effect of group or interaction of the group with other factors. We noted that the greater variability around zero PTE in the group of amusic participants, for delta-band connectivity between the auditory to the inferior frontal cortices, and between the auditory and the motor cortices, did not indicate clear directionality during tone sequence presentation (both during pre- and post-target segments, t-test against zero p > 0.099).

Akin to typical listeners, top-down beta-range directed transfer was significant from motor to auditory cortices (resting-state: t(15) = −6.49, p < 0.001, pre: t(15) = −7.67, p < 0.001, post: t(15) = −8.11, p < 0.001), and from motor to inferior frontal cortices (resting-state: t(15) = −6.31, p = 0.005, pre: t(15) = −6.80, p < 0.001, post: t(15) = −4.44, p = 0.009). There was no difference between the right and left hemisphere in amusics (F < 1.33, p > 0.25).

## Discussion

We used non-invasive neurophysiological measures of local and inter-regional brain dynamics to study the neurophysiological mechanisms of pitch change detection. These analyses were performed in typical listeners and in congenital amusics, to resolve the mechanistic elements that are essential to pitch perception, and deficient in amusia.

Our behavioral results confirmed that amusics had more difficulty than controls in detecting small pitch variations of up to 50 cents (Fig. 1B). We show in Supplementary Material (Supplementary Fig. S2) that lower accuracy of amusic participants was not related to lack of vigilance while performing a task that was difficult and possibly frustrating to them (Ciesielski et al., 2007). Indeed, we found that posterior alpha-band activity – as a proxy marker of decreased vigilance – was lower in amusics than in controls during the task (Valentino et al., 1993; Aftanas and Golocheikine, 2001). This observation could be attributed to amusics experiencing the task as more difficult hence soliciting more vigilance from their part, marked with lesser alpha power.

Our data report effects within and between brain regions of interest that have been repeatedly highlighted in previous pitch processing studies using a variety of functional techniques (Zatorre et al., 1992; Albouy et al., 2013; Peretz, 2016; Morillon and Baillet, 2017). These regions comprise, bilaterally, the superior temporal gyrus in auditory cortices, the posterior portion of the inferior frontal gyrus and the pre-central motor cortices.

During the presentation of tone sequences, we found local expressions of cross-frequency coupling between the phase of delta-band activity and the amplitude of beta-band signal components in the right auditory and inferior frontal cortices in both groups. The frequency of delta-band activity was similar to the presentation rate of tones in the auditory sequences (2.85 Hz), which is typical of cortical entrainment at the dominant rate of auditory signals (Doelling and Poeppel, 2015; Morillon and Baillet, 2017, Puschmann et al. 2018). By boosting neural signals in response to regular sensory inputs, cortical entrainment increases signal-to-noise ratio and improves the detection of genuine phase-amplitude coupling effects (Aru et al., 2015; Samiee & Baillet, 2017). There was no delta-to-beta coupling above chance level in the absence of tone-sequence presentation, namely during baseline resting state in IFG (Fig. 2G). The fact that we observed delta-band entrainment in IFG with the task (Fig. 2B-C) is compatible with this region being a downstream node of the ventral auditory pathway (Zatorre et al., 1992; Gaab et al., 2003, Albouy et al., 2013, 2019). Expressions of beta-band activity during pitch processing have been previously reported in auditory regions (Cirelli et al., 2014; Fujioka et al., 2012), including during the pre-target time period (Florin et al., 2017).

In both groups, delta-beta phase-amplitude coupling was elevated in auditory and inferior frontal cortices during task performance compared to baseline resting state (Fig. 2G). This observation is in line with reports of higher transient PAC levels during task performance, such as with working memory (Axmacher et al., 2010), associative learning (Tort et al., 2009; van Wingerden et al., 2014) and visual attention (Szczepanski et al., 2014).

A striking overall effect between groups was that delta-to-beta coupling in the auditory and inferior frontal cortices was higher in amusics than in controls, both during tone-sequence presentations and at baseline in the resting state. These observations of elevated ongoing phase-amplitude coupling are the first observed in amusia. They contribute to converging evidence that chronically elevated PAC levels could be brain signal indicators of impaired neurophysiological function, as previously shown in e.g., epilepsy (Amiri et al., 2016; Samiee et al., 2018), Parkinson’s disease (van Wijk et al., 2016) and autism spectrum disorders (Berman et al., 2015). In amusic participants, we found increased PAC levels in regions that have been reported as abnormal in congenital amusia using structural MRI (Hyde 2006, Albouy et al., 2013), malfunctioning with functional MRI (Hyde et al., 2011, Albouy et al., 2019) and electrophysiology (Albouy et al., 2013, 2015, Tillmann et al., 2016). Our perspective is that this observation is compatible with previous reports of stronger expressions of slow (delta-range) prediction error signaling in the auditory cortex of amusics, during presentation of tone sequences (Albouy et al., 2015). Recent data on neurophysiological processing of natural speech also reported that delta signaling is enhanced in auditory cortex by words and phonemes that are less predictable in the sentence flow (Donhauser & Baillet, 2020). Mechanistically, we propose that although PAC is expressed ubiquitously and dynamically in the human brain (Florin & Baillet, 2015), over-expressions of PAC coupling may reflect a lack of flexibility in the adjustment of the phase angle where fast frequency bursts are nested along slow frequency cycles. This phase angle is related to the level of net excitability of the underlying cell assemblies and has been discussed as an essential parameter for the neural registration of sensory inputs (Gips et al., 2016). High levels of PAC may reduce opportunities for registering, and therefore encoding and processing, incoming sensory inputs with sufficient temporal flexibility and adaptation to prediction errors (Arnal & Giraud, 2012). These considerations may inspire future studies in the field.

Delta-to-beta coupling was stronger in auditory regions than in inferior frontal cortex in both groups (Fig. 2G), which we interpret as due to the entrainment of auditory delta activity by stimulus cortical inputs, which is expected to be more direct than in downstream regions. Yet, another marked difference between groups was that there were modulations of delta-to-beta phase-amplitude coupling in the inferior frontal cortex of typical listeners depending on their individual report of the target tone being perceived as deviant, regardless of accuracy. Such percept-dependent increases are compatible with the known involvement of the inferior frontal cortex in pitch detection (Doeller et al., 2003; Alain et al., 2001; Florin et al., 2017) and in integrating auditory events that are presented sequentially (Tillmann et al. 2006, Albouy et al., 2013, 2017). There was no such modulation in amusics, which is in line with the absence of P300 event-related responses in this population for small pitch deviations (see also Supplementary Fig.S1), with IFG as a contributing generator (Florin et al., 2017; Albouy et al. 2013; 2015).

We derived time-resolved measurements of phase-amplitude coupling (tPAC) over time windows around the occurrence of each of the tones in the sequence. In the auditory cortex of both groups, there was an increase of cross-frequency coupling immediately after the onset of the tone sequence (Fig. 2B), which culminated at the expected latency of the target tone presentation. This was confirmed by a time - resolved analysis of stimulus-to-beta coupling in the auditory cortex, which showed that stronger phasic beta activity occurred at the expected latency of the auditory tones in the sequence (Fig. 2H). This observation is compatible with the signaling of predictive inferences concerning the timing of the next expected tone presentation in the sequence, which Morillon and Baillet (2017) showed to be emphasized by temporal attention. These effects were observed both in typical listeners and amusics. Such coupling between the timing of tone presentations (i.e. the actual physical stimulus) and modulations of beta-band signal amplitude in auditory cortex was previously observed by Chang et al. (2018): They showed that stimulus-to-beta coupling in right auditory cortex was associated with the predictability of pitch changes in a sequence. In our study, pitch changes occurred systematically on the fourth tone of the sequence and in only 50% of the trials. Hence, the predictability of the timing of pitch changes was high but their actual occurrences were poorly predictable from trial to trial. In that respect, our results are consistent with Chang et al. (2018)’s, as our participants presented lower levels of stimulus-to-beta coupling than endogenous delta-to-beta, with no temporal modulations along the presentation of the tone sequence (Fig. 2H).

We interpret the lesser levels of stimulus-to-beta compared to delta-to-beta coupling as due to the fact that the dominant delta-band neurophysiological activity did not exactly match the tone presentation rate. This is indicative of phase and frequency jitters between the regular auditory inputs of the tone sequence and the induced neurophysiological responses. We also observed (Fig. 2G) that the beta-band activity in auditory cortex was modulated by target-tone presentations more strongly in amusics than in controls. At the present time, we can only speculate that this may reflect the allocation of greater neural computation resources locally in primary auditory regions for tone prediction in amusics.

Our observations of functional connectivity, at different rhythmic frequencies of neural activity, between regions of interest provide further insight into both the neurophysiological processes of typical pitch change detection and of the impaired processing in amusia. Coherence measures revealed stronger delta-band effects, right-hemisphere dominance and auditory-IFG connectivity in typical listeners than in amusics, confirming previous published observations (Albouy et al., 2013; 2015; 2019; Peretz, 2016).

Directed connectivity analyses during resting-state baseline revealed bilateral influences in the beta band from the motor cortex, directed to the auditory and inferior frontal cortices. These interactions persisted during task performance and were emphasized during the pre-target segment of each trial. These results are in line with reports of dynamically structured and anatomically organized beta-band activity in the resting state (Brookes et al., 2011; Bressler and Richter, 2015). They are also concordant with strong emerging evidence that beta-band activity is a vehicle for top-down signaling in brain systems during sensory processing (Engel et al., 2001; Engel & Fries, 2010; Bressler & Richter, 2015; Bastos et al., 2015; Michalareas et al., 2016; Morillon & Baillet, 2017; Chao et al., 2018). This body of empirical evidence is in support of the theoretical framework of predictive coding (Rao & Ballard, 1999; Friston & Kiebel, 2009) and predictive timing (Arnal & Giraud, 2012; Morillon & Baillet, 2017) in sensory perception. In this context, beta-band top-down activity would channel predictive information concerning the expected nature and temporal occurrence of incoming sensory information to primary systems (Fontolan et al., 2014; Baillet, 2017). In essence, the theoretical principles posit sensory perception as an active sensing process, in which the motor system would play a key role especially in predicting the timing of expected sensory events (Schroeder et al., 2010). In audition for instance, we previously showed that, even in the absence of overt movements, beta-band oscillations issued in the motor cortex had influence on auditory cortices and contributed to the temporal prediction of tone occurrences in complex auditory sequences (Morillon and Baillet, 2017). Our present data confirm and extend these observations: the modulations of beta-band activity in the auditory cortex peaked at the expected and effective occurrences of the tones in the sequence, which is compatible with the involvement of the motor cortex in driving inter-regional signals for predictive sensory timing. This top-down signaling mechanism was not affected in amusic participants.

During tone-sequence presentations, we found only in controls a bottom-up form of directed connectivity issued from the auditory cortices towards both the inferior frontal and motor cortices (Fig. 3B). These connections were not significantly expressed in amusics and were not present during the baseline resting state in both groups. Delta-band oscillatory activity contributed to the mode of maximum regional phase-amplitude coupling and encompassed the stimulus presentation rate of 2.85 Hz. Coherence connectivity analysis reflected that there was a hemispheric asymmetry connectivity transfer towards the right hemisphere (Fig. 3A), in line with previous reports (Zatorre et al., 1992). Such bottom-up connectivity transfer is also compatible with the principles of predictive coding and timing, which posit that primary sensory regions propagate prediction-error signals downstream in brain systems networks, for ongoing updates of internal predictive and decision models (Rao & Ballard, 1999; Friston & Kiebel, 2009; Baillet, 2017) as recently shown in natural speech processing (Donhauser & Baillet, 2020). This observation is consistent with published dynamical causal models of impaired directed connections between auditory and inferior frontal cortices in amusics (Albouy et al., 2013; 2015) and other neurophysiological disorders (Omigie et al., 2013). These previous results were not specific to narrowband oscillatory signals: they were obtained from event-related signals in response to tone-sequence presentations. Our findings are also in line with fMRI data showing reduced functional – not directed – connectivity in amusic participants between the same ROIs (Hyde et al., 2011, Albouy et al., 2019). Loui et al. (2009) also reported reduced anatomical connections via the arcuate fasciculus in amusic participants using diffusion-weighted imaging and tractography, although more recent results have been mixed (Chen et al., 2015; Wilbiks et al., 2016). In our data, directed connectivity measures were qualitatively similar between amusics and controls. The inter-individual variability of directed connectivity statistics was greater in amusics, which may explain why both the strength and directionality of connections were deemed not significant in this group. We acknowledge that our sample size was small – due to the relatively rare amusia syndrome. Yet the motifs of directed connectivity are compatible with the large effects we observed in behavior, local phase-amplitude coupling statistics, and functional connectivity statistics (coherence) reflecting stronger delta-band interactions in typical listeners than in amusics.

In conclusion, we provide evidence that pitch discrimination from a sequence of pure tones engages a distributed network of cortical regions comprising at least the auditory, inferior frontal and lateral motor cortices. We also show that the motor cortex issues beta-band signals directed to inferior frontal and auditory regions, which are present by default in the resting state, but which timing during auditory presentation marks the actual expected occurrences of tones in the sequence. The auditory cortex is entrained at a rate around the physical pace of the tone sequence, and this signal is propagated in a bottom-up fashion further downstream to the motor system and along the ventral pathway to the inferior frontal cortex. These poly-frequency phenomena interact locally through phase-amplitude coupling, which increases in auditory regions at the onset of the tone sequence and culminates at the expected occurrence of the target tone before returning to baseline levels. Our data identify two cross-frequency mechanisms as crucial to pitch-change detection, when contrasting amusic participants with typical listeners. First, delta-to-beta phase-amplitude coupling is chronically elevated in the auditory and inferior frontal regions of amusics. Second, bottom-up signaling along the ventral auditory pathway and to the motor cortex is depressed in this group. In sum, our findings point at an alteration of pitch encoding in the auditory regions of amusics, which may depress prediction error signaling driven to motor and inferior frontal regions, and eventually poorer perceptual detection. The predictive timing functions seem to be preserved in amusics, at least in the present context of highly predictable and regular pacing of the tone sequence.

Taken together, we believe these findings advance the complete and dynamic view of tone sequence sensory processing in audition. We anticipate that some of these new observations would generalize to other sensory modalities and that the cross- and poly-frequency neurophysiological markers of impaired auditory processing would be pertinent to other functional deficits in sensory perception.

## Material and Methods

### Participants

Sixteen right-handed participants took part in the study. Eight amusics were recruited according to the Montreal Protocol for Identification of Amusia (MPIA; Vuvan et al., 2018). The other eight participants formed a control group (i.e. typical listeners) who were matched with the amusic group in terms of age, gender, and years of education. The experimental paradigm was reviewed and approved by the ethics review board of McGill University Health Centre (Protocol: NEU-12-023). All participants gave written informed consent to take part in the study.

All participants had their hearing assessed using age-normalized audiometric guidelines (Park et al., 2016). Although some participants had heightened audiometric thresholds (which is common in older adults), all participants reported being able to hear the experimental trials without effort. Supplementary Table S1 displays demographic information for all participants and reports their performances on a battery of musical tasks. The amusic and control participants were matched in terms of demographic variables but differed significantly in musical performance. Specifically, amusics scored significantly lower than controls on the melodic tasks (AMUSIA Scale, AMUSIA Out-of-Key, MBEA Melodic). In contrast, they scored equivalently to controls on the AMUSIA Off-Beat test, confirming the specificity of their disorder to musical pitch.

### Experimental design

The study paradigm was adapted from Hyde and Peretz (2004) and Peretz et al. (2005). Each trial consisted of a sequence of five pure tones: Tones #1, 2, 3 and 5 were identical and played at the pitch level of C6 (1047 Hz; standard pitch). Tone #4 was the target tone played at 5 different pitches across trials. In half of the trials, the target tone was played at the standard C6 (1047 Hz) pitch (“standard” trials). In the other half of the trials (“deviant” trials), the target tone was played with a deviation of 25, 50, 100, or 200 cents (100 cents correspond to 1 semitone) from the standard tone. Each tone was presented for 100 ms, and the time interval between two consecutive tone onsets in a sequence (inter-tone interval, ITI) was 350 ms. The total duration of a sequence was 1.4 s (Fig. 1A).

Ten minutes of resting-state were recorded from all participants (eyes open) at the beginning of the session. Participants were then asked to listen to tone sequences and to press a button with one of their index fingers to indicate whether the presented sequence comprised a standard or a deviant target sound. They were instructed to keep their gaze fixed on a cross displayed on a back-projection screen positioned at a comfortable distance. Responses with the right- or left-hand finger to standard vs. deviant trials were intermixed between participants. All subjects received 40 training trials prior to MEG data collection. A total of 640 tone sequences were then presented to every participant, in 10 blocks of 64 trials, which resulted in a total of 320 standard tone sequences and 80 deviant trials per pitch deviance level. Trials started in succession, 1 second (± < 50 ms jitter) following the subject’s response to the previous trial. No feedback was provided to participants on their performance.

### Data acquisition

MEG data was collected during resting-state and task performance in the upright position using a 275-channel CTF MEG system, with a sampling rate of 2400 Hz. Simultaneous EEG data was recorded also using the CTF system from four standard electrode positions: FZ, FCZ, PZ, and CZ (reference was placed on right mastoid), electrode locations according to 10/20 system with 2400 Hz sampling rate (EEG data shown as Supplementary Material – Fig. S1). The audio presentation, button presses, heartbeat and eye movement electrophysiological signals (ECG and EOG, respectively) were also collected in synchronization with MEG. Head position was monitored and controlled using three coils attached to the subject’s nasion). We obtained T1-weighted MRI volumes for each participant (1.5-T Siemens Sonata, 240 × 240 mm field of view, 1 mm isotropic, sagittal orientation) for cortically-constrained MEG source imaging (Baillet, 2017). Polhemus 3-D digitizer system (https://polhemus.com/scanning-digitizing/digitizing-products/). We obtained T1-weighted MRI volumes for each participant (1.5-T Siemens Sonata, 240 × 240 mm field of view, 1 mm isotropic, sagittal orientation) for cortically-constrained MEG source imaging (Baillet, 2017).

### Data preprocessing and source modeling

Contamination from system and environmental noise was attenuated using built-in CTF’s 3rd-order gradient compensation. All further data preprocessing and modeling was performed with Brainstorm (Tadel et al., 2011) following good-practice guidelines (Gross et al., 2013). The recordings were visually inspected, with segments contaminated by excessive muscle artifacts, head movements or remaining environmental noise marked as bad and discarded from further analysis. Powerline artifacts at 60 Hz and harmonics up to 240 Hz were reduced using notch filtering. Signal-space projectors (SSP) were designed using Brainstorm’s default settings to attenuate the electrophysiological contamination from heartbeats and eye blinks.

The MRI data was segmented using the default FreeSurfer pipeline (Dale et al. 1999). For distributed source imaging, we used Brainstorm to resample the cortical surface tessellation produced down to 15,000 vertices. We derived individual forward MEG head models using the overlapping-sphere analytical approach (with Brainstorm default settings). We then obtained a weighted minimum-norm kernel (wMNE; Brainstorm with default settings) for each participant to project sensor-level preprocessed data onto the 15,000 vertices of the individual cortical surface. The empirical covariance of sensor noise was estimated for wMNE modeling from a 2-min empty-room MEG recording collected at the beginning of each session, i.e. for each participant.

### Regions of interest

In all participants, we defined six brain regions of interest (ROIs) using a MEG functional localizer. The right and left auditory cortices (rAud and lAud) were identified as the regions presenting the strongest M100 (within 100-120 ms post-stimulus onset) event-related average response peak to all tones, restricted to 3cm^2^ of surface area per region. We defined rIFG and lIFG as portions of Brodmann BA45 identified from the Brodmann cortical atlas of Freesurfer registered to individual anatomy. The spatial extent of the rIFG and lIFG ROIs was based on the maximum differential activity observed between the brain responses to deviant and standard tones around 100 ms after “target tone” presentation. The resulting surface areas varied between ROIs as driven by the strength of the event-related response observed and were typically about 1.3 cm^2^. Left and right cortical motor regions (lMot and rMot, respectively) were defined following Morillon and Baillet (2017) over a surface area of about 3cm^2^ at the pre-central locations of the largest M50 50-ms latency responses after right and left index finger button presses, respectively (Fig. 1D).

### Phase-amplitude coupling

We used the time-resolved measure of phase-amplitude coupling (tPAC) between co-localized slow and fast cortical signal components, as published by Samiee and Baillet (2017) and distributed with Brainstorm. tPAC measures the temporal fluctuations of the coupling between the phase of slower activity (at frequency *fP*) and the amplitude of faster signal components (at frequency *fA*). Briefly, the instantaneous amplitude of faster signals (A_fA_ (t)) in a sub-band of the *fA* band of interest was extracted using the Hilbert transform. Power spectral analysis was used to identify the frequency of strongest oscillation in *A*_*fA*_ (*t*) (in the *fP* band of interest), coinciding with an oscillation in the original time series. This frequency was then labeled as the *fP* frequency coupled to the current fast *fA* frequency. The coupling strength between *A*_*fA*_ (*t*) and the instantaneous phase of the signal filtered around *fP* were then calculated. See Samiee and Baillet (2017) for further methodological details concerning tPAC.

We then extracted comodulograms to identify the strongest modes of (*fP*, *fA*) coupling over time windows of 1.5 s that contained the entire tone sequence at every trial, testing 20 candidate *fA* frequencies linearly distributed within the [15, 250]-Hz band. The frequency band of interest for *fP* was [2, 12] Hz. The strongest (*fP, fA*) mode of coupling was identified from the obtained comodulogram with *fP* in the [2, 4] Hz band and *fA* in the [15, 35] Hz band. The temporal dynamics of tPAC coupling were extracted between these two bands of interest (reflecting the dominant mode of coupling) from 700-ms time windows with 50% overlap over the entire trial duration. Since previous studies reported on right-hemisphere dominance in similar pitch discrimination tasks (Peretz, 2016; Zatorre et al., 1992), we analyzed phase-amplitude coupling in the right-hemisphere ROIs only.

### Stimulus-brain coupling

We assessed whether the auditory stimulus tone sequence induced modulations of beta activity in the auditory cortex. The goal was to replicate previous observations of stimulus-induced beta-amplitude modulations in auditory cortex in similar conditions (Fujioka et al., 2012; Cirelli et al., 2014; Chang et al., 2018). These results emphasize how beta-band activity is expressed by the auditory tone sequence instead of local delta activity in the auditory cortex, and whether beta bursts occur preferentially at the expected latency of the tone presentation as a predictive form of signal.

Following the method used by Morillon and Baillet (2017), we generated a reference sinusoidal signal at 2.85 Hz (i.e. the rate of the tone presentation every 350 ms), with its peaks aligned at the onset of each tone presentation. We then estimated the tPAC cross-frequency coupling between the phase of this reference signal and the amplitude of beta oscillations in the right auditory cortex. We tracked the variations in time of this coupling using tPAC with a sliding window length of two cycles of the tone presentation rate (700 ms) with 50% of overlap, following the specifications derived by Samiee and Baillet (2017). We then identified the preferred phase of tPAC coupling along the cycle of the stimulus sinusoid reference signal. Finally, we converted the corresponding phase angle into a time latency, as a fraction of the 350 ms stimulus presentation cycle.

### Functional and effective connectivity

We estimated frequency-specific functional connectivity between ROIs using coherence (Walter et al., 1966; Thatcher et al. 1986; Fries, 2005), which is a measure of amplitude and phase consistency between cortical signals. We also measured signs of directional, effective connectivity between ROIs with phase-transfer entropy (PTE; Lobier et al., 2014), adopting the approach by Morillon and Baillet (2017). PTE measures narrowband phase leading/lagging statistics to derive estimates of effective connectivity between regions. Importantly to our study, PTE has shown better performance than coherence in detecting signs of interdependence between signals of relatively short duration (Bowyer et al., 2016). In more details, PTE measures effective connectivity based on the respective instantaneous phases of pairs of narrow-band neurophysiological signals. The sign of directed PTE (dPTE) values indicate the estimated direction for effective connectivity. For example, considering two regions A and B, positive (respectively negative) dPTE values indicate information transfer from A (resp. B) to B (resp. A). We used the dPTE code openly shared by Hillebrand et al. (2016), which we have made available in Brainstorm.

Informed by tPAC frequency ranges, we derived dPTE directed connectivity measurements between the regions of interest and between hemispheres, in the frequency bands of interest. We evaluated interhemispheric coherence and dPTE connectivity between homologous regions bilaterally, in the delta ([2,4] Hz) and beta ([15,30] Hz) frequency bands, over the baseline resting-state period, the [−1500,0] ms pre-target and the post-target [0,1500] ms time segments.

In reported results, the significance of observed dPTE values in each group and condition was statistically corrected for all comparisons performed (18 comparisons for: 3 pairs of regions × 2 frequency bands × 3 states).

### Statistical analyses

Parametric tests (e.g., t-tests against zero-mean, paired t-tests, repeated measures ANOVAs) were employed (with p = 0.05 considered as significance threshold). Tukey’s tests were used for post-hoc analyses and corrections for multiple comparisons. The distributions of event-related potentials (see Supplementary Material) were tested for zero-mean using t-tests and reported with corrections for multiple comparisons considering false discovery rates (FDR). tPAC values were assessed for statistical significance using a non-parametric resampling approach (Samiee and Baillet, 2017): for each trial, we generated 500 surrogates using block-resampling. Each surrogate was produced from selecting five time points randomly in the trial epoch to subdivide the instantaneous phase signal into five blocks. These blocks were then randomly shuffled and tPAC was estimated using the resulting block-shuffled phase signal and from the original instantaneous amplitude time series. This resampling technique provides reference surrogate signals with phase-amplitude coupling at chance levels and with minimum phase distortion (Samiee and Baillet, 2017). The tPAC values obtained from surrogate data were normally distributed (Shapiro-Wilk test, p > 0.8). tPAC values from each original trial were z-scored with respect to the empirical distribution of tPAC values obtained from the surrogate data generated fr om the same trial

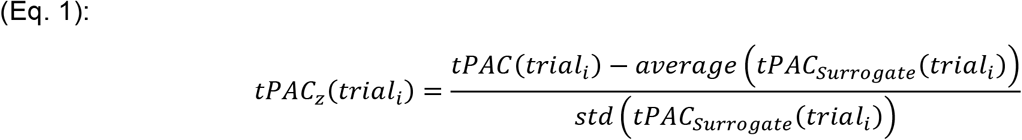

## Acknowledgment

S.S. acknowledges the support from McGill University Integrated Program in Neuroscience. I.P. was supported by grants from the Natural Sciences and Engineering Research Council of Canada, the Canadian Institutes of Health Research, and the Canada Research Chair program. S.B. was supported by a Discovery grant from the Natural Science and Engineering Research Council of Canada (436355-13), the Canada Research Chair of Neural Dynamics of Brain Systems, the NIH (1R01EB026299-01), the Healthy Brains for Healthy Lives Canada Excellence Research Fund and a Platform Support Grant from the Brain Canada Foundation (PSG15-3755). Pilot data for this study was collected with support from Research Incubator Grant from McGill University’s Center for Research on Brain Language & Music.

## Author Contributions

SS was responsible for designing the analysis methods, writing the corresponding codes; she performed most data analyses, produced all figures, and was the primary author of all drafts and the final version of the manuscript. EF and DV ran the data collection and were involved in the study design. PA contributed to the analysis approach and reviewed the final version of the manuscript. IP was responsible for the study design, co-funding, and reviewed the final version of the manuscript. SB was responsible for supervising the project, co-funding, as well as editing drafts and the final version of the manuscript.

