## Supplementary material for "Neurophysiological network dynamics of pitch change detection"

**This PDF file includes:**

Supplementary text
Tables S1
Figures S1 to S2
SI References

### Participants

Details of participants demographics is shown in the following table (Table S1).

Supplementary Table S1. Participant demographics and musical test performances. The data are reported as group mean and standard deviation [SD]. For a detailed description of the musical tests employed, see (Vuvan et al., 2018).

| | | AMUSICS | CONTROLS | STATISTICAL TEST |
|---|---|---|---|---|
| **DEMOGRAPHICS** | N | 8 | 8 | --- |
| | # women | 6 | 6 | --- |
| | age (years) | 65.13 [6.10] | 63.14 [5.96] | $t(14) = 0.51$, $p = .62$ |
| | education (years) | 16.13 [2.17] | 15.43 [2.76] | $t(14) = 0.89$, $p = .39$ |
| | music education (years) | 1.00 [1.07] | 1.57 [1.51] | $t(14) = 1.16$, $p = .27$ |
| **MUSICAL TESTS** | AMUSIA Scale (/100) | 55.42 [10.15] | 91.25 [6.56] | $t(14) = 8.39$, $p < .001$ |
| | AMUSIA Off-Beat (/100) | 70.84 [10.79] | 78.65 [6.05] | $t(14) = 1.79$, $p = .09$ |
| | AMUSIA Out-of-Key (/100) | 54.29 [10.62] | 84.56 [7.83] | $t(14) = 6.53$, $p < .001$ |
| | MBEA Melodic (/30) | 18.25 [2.63] | 26.71 [2.60] | $t(13) = 6.47$, $p < .001$ |

### Event-related responses

**Method**

Event-related brain responses were extracted from the time-locked average of correctly detected trials in all experimental conditions (standard and various degrees of pitch deviance). Electrophysiological signals were bandpass filtered between [0.5, 50] Hz and baseline corrected with respect to the 150 ms before the presentation of the target tone ([-150, 0] ms).

**Results**

Group average event-related responses to target tone presentations are shown for electrode CZ in Fig. S1. There was a clear N1 component around 110 ms following the onset of the target tone in both groups and all three conditions. In line with previous reports (Peretz et al., 2005), both groups produced a P3 component in the high-deviance condition (significantly different from zero-mean $t(7) > 4.8$, $p < 0.002$). The P3 response was weaker for standard tones in both groups. In the low-deviance condition, amusics showed similar responses than to standards (not significantly different from zero-mean $t(7) < 2.3$, pF DR > 0.05), with controls producing a P3 which amplitude was intermediate between those in standard and high-deviance conditions (significantly different from zero-mean $t(7) > 3.5$, $p < 0.01$).

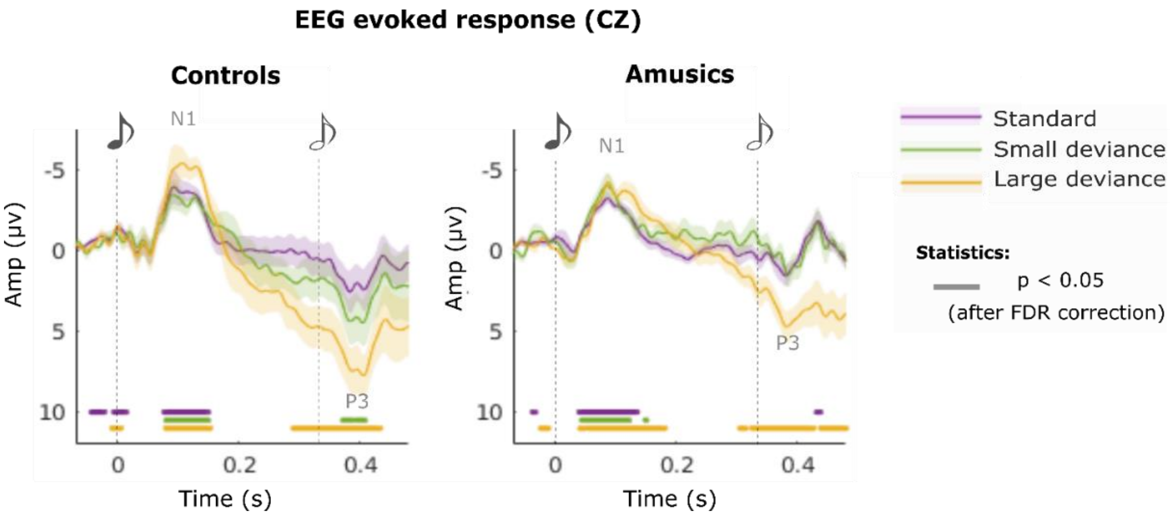


**Supplementary Figure S1:** EEG event-related responses in the standard, small and large deviance conditions in both groups (CZ electrode).

### Monitoring of vigilance: posterior alpha-band activity

A cluster of five posterior MEG channels presenting the highest levels of pre-target [8, 12]-Hz alpha-band activity across subjects (MZP01, MLP31, MRP31, MLP32, MRP32) were selected. The power of MEG signals at these sensor locations over the pre-target period ([-1.5, 0] s) of each trial was computed. We used the even-order linear-phase FIR filter in Brainstorm (band-pass: [8, 12] Hz, stop-band attenuation: 40 dB, 99% energy transient: 0.402 s) and computed the root-mean-square (RMS) signal strength across the sensor cluster for each trial. The same approach was applied to EEG, restricted to electrode PZ.

We measured the posterior normalized alpha power as a proxy for vigilance (Valentino et al., 1993), attention (Aftanas & Golocheikine, 2001) and the cognitive demand of the task (Gevins & Smith, 2000; Ciesielski et al., 2007). Higher alpha levels could indeed account for lower task performances and confound the interpretation of our data. There was a main effect for group and task performance (Supplementary Fig. S2). Amusics produced lower levels of posterior alpha activity ($F(1)=28.37$, $p < 0.001$), which could be indicative of the task requiring higher attentional demands from this group (Gevins et al., 1979; Smith et al., 1999). Posterior alpha activity was reduced in correct trials ($F(1)=8.1$, $p = 0.004$), which is consistent with its negative association with attention and vigilance. Interactions between response accuracy and groups were significant ($F(1)=5.86$, $p = 0.01$), with a post-hoc Tukey's test showing lower alpha power in correct trials in amusics ($t(9262)=-4.46$, $p_{Tukey} < 0.001$) and significantly lower posterior alpha levels in amusics compared to controls in both correct and incorrect trials (correct: $t(9262)=3.98$, $p_{Tukey} < 0.001$, incorrect: $t(9262)=4.16$, $p_{Tukey} < 0.001$). We made similar observations from EEG recordings at electrode Pz (not shown).

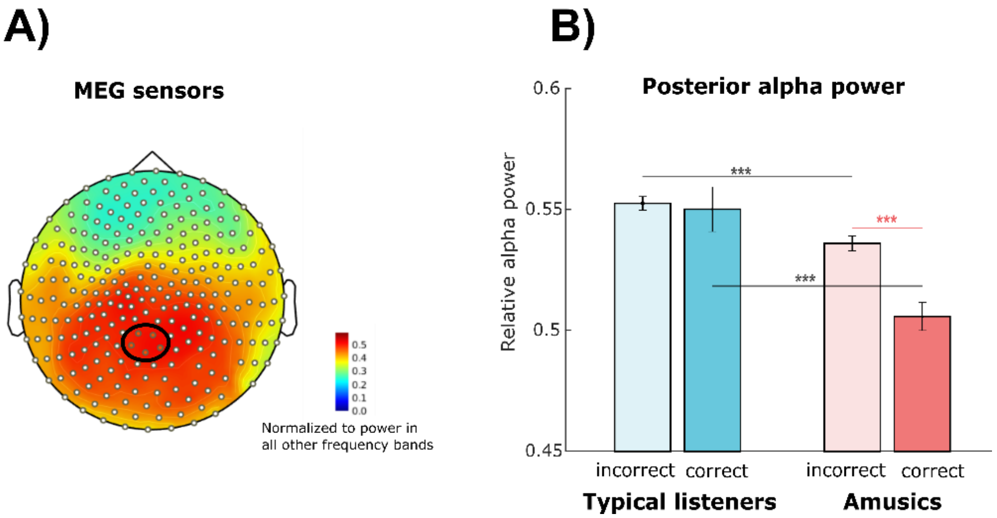


**Supplementary Figure S2**: Posterior alpha power averaged over trial baselines (pre-target period) across a cluster of posterior MEG sensors as a measure of vigilance. A) Normalized alpha power (to the power of all other frequency bands) in a representative subject, reflecting the selected MEG sensors for alpha power measurement (in black oval) .B) Average MEG posterior alpha power: interaction of groups and correctness of trials.
